# Saturation genome editing-based functional evaluation and clinical classification of BRCA2 single nucleotide variants

**DOI:** 10.1101/2023.12.14.571597

**Authors:** Huaizhi Huang, Chunling Hu, Jie Na, Steven N. Hart, Rohan David Gnanaolivu, Mohamed Abozaid, Tara Rao, Yohannes A. Tecleab, Tina Pesaran, Paulo Cilas Morais Lyra, Rachid Karam, Siddhartha Yadav, Susan M. Domchek, Miguel de la Hoya, Mark Robson, Miika Mehine, Chaitanya Bandlamudi, Diana Mandelker, Alvaro N.A. Monteiro, Nicholas Boddicker, Wenan Chen, Marcy E. Richardson, Fergus J. Couch

## Abstract

Germline BRCA2 loss-of function (LOF) variants identified by clinical genetic testing predispose to breast, ovarian, prostate and pancreatic cancer. However, variants of uncertain significance (VUS) (n>4000) limit the clinical use of testing results. Thus, there is an urgent need for functional characterization and clinical classification of all *BRCA2* variants. Here we report on comprehensive saturation genome editing-based functional characterization of 97% of all possible single nucleotide variants (SNVs) in the BRCA2 DNA Binding Domain hotspot for pathogenic missense variants that is encoded by exons 15 to 26. The assay was based on deep sequence analysis of surviving endogenously targeted haploid cells. A total of 7013 SNVs were characterized as functionally abnormal (n=955), intermediate/uncertain, or functionally normal (n=5224) based on 95% agreement with ClinVar known pathogenic and benign standards.

Results were validated relative to batches of nonsense and synonymous variants and variants evaluated using a homology directed repair (HDR) functional assay. Breast cancer case-control association studies showed that pooled SNVs encoding functionally abnormal missense variants were associated with increased risk of breast cancer (odds ratio (OR) 3.89, 95%CI: 2.77-5.51). In addition, 86% of tumors associated with abnormal missense SNVs displayed loss of heterozygosity (LOH), whereas 26% of tumors with normal variants had LOH. The functional data were added to other sources of information in a ClinGen/ACMG/AMP-like model and 700 functionally abnormal SNVs, including 220 missense SNVs, were classified as pathogenic or likely pathogenic, while 4862 functionally normal SNVs, including 3084 missense SNVs, were classified as benign or likely benign. These classified variants can now be used for risk assessment and clinical care of variant carriers and the remaining functional scores can be used directly for clinical classification and interpretation of many additional variants.

**Summary:** Germline BRCA2 loss-of function (LOF) variants identified by clinical genetic testing predispose to several types of cancer. However, variants of uncertain significance (VUS) limit the clinical use of testing results. Thus, there is an urgent need for functional characterization and clinical classification of all *BRCA2* variants to facilitate current and future clinical management of individuals with these variants. Here we show the results from a saturation genome editing (SGE) and functional analysis of all possible single nucleotide variants (SNVs) from exons 15 to 26 that encode the BRCA2 DNA Binding Domain hotspot for pathogenic missense variants. The assay was based on deep sequence analysis of surviving endogenously targeted human haploid HAP1 cells. The assay was calibrated relative to ClinVar known pathogenic and benign missense standards and 95% prevalence thresholds for functionally abnormal and normal variants were identified. Thresholds were validated based on nonsense and synonymous variants. SNVs encoding functionally abnormal missense variants were associated with increased risks of breast and ovarian cancer. The functional assay results were integrated into a ClinGen/ACMG/AMP-like model for clinical classification of the majority of BRCA2 SNVs as pathogenic/likely pathogenic or benign/likely benign. The classified variants can be used for improved clinical management of variant carriers.

## Introduction

*BRCA2*, an established clinically actionable cancer predisposition gene^1^, has been widely used for hereditary cancer genetic testing. The knowledge obtained from testing has been effectively used to assess risks of various cancers, with BRCA2 pathogenic variants associated with 69% lifetime breast cancer risk^2^ and 15% ovarian cancer risk^3^ and substantially increased risk of pancreatic and prostate cancer^4,5^. Genetic testing and associated pathogenic variants are now effectively used for clinical management of carriers, in terms of prevention, screening and cancer treatment. Broader acceptance and application of genetic testing in recent years has resulted in the identification of large numbers of individual variants. However, the interpretation and classification of the majority of these variants, despite detailed review of multiple sources of genetic and clinical information, has not been possible mainly due to rarity in the general population. Over 4000 individual *BRCA2* variants are currently classified on ClinVar^6^ as variants of uncertain significance (VUS). These unclassified variants, that are largely missense and intronic alterations, cannot be effectively utilized for clinical care. Thus, there is an urgent need for large scale characterization and classification of *BRCA2* variants.

Previously, multifactorial prediction models using a prior probability of pathogenicity, personal or family history, and co-segregation of variants with cancer, was used to classify a small number of VUS in BRCA2 as pathogenic or benign^7–9^. Recently, American College of Medical Genetics (ACMG)/Association for Molecular Pathology (AMP) guidelines that incorporate multiple sources of evidence including family history, variant frequency in populations, *in silico* sequence-based prediction, functional data, and others, have been utilized by clinical testing groups and the ClinGen BRCA2 variant curation expert panel (VCEP) for classification of variants^10^. Due to limited information from other types of evidence, classification using these models is heavily dependent on functional assay results for classification of variants. Previous work integrated functional data for BRCA2 missense variants from a well validated and risk calibrated homology directed repair (HDR) assay into an ACMG/AMP framework resulting in classification of more than 400 BRCA2 DNA Binding Domain (DBD) missense variants as pathogenic or benign^11–13^.

However, this and other functional assays have been low-throughput and have not substantially resolved the VUS issue. In contrast, multiplex assays of variant effect (MAVEs) allow for the parallel functional characterization of large numbers of variants^14^. Using cell-based or *in vitro* selection and deep sequencing to link genotype to phenotype, many variants can be functionally characterized and compared to results for known pathogenic and benign standards. These methods have been effectively applied to functional characterization of SNVs in cancer predisposition genes including *BRCA1* and *MSH2*^15,16^. MAVE analysis of *BRCA2* has been predominantly in the form of proof of principle studies that focused on relatively small regions of *BRCA2* ^17–19^, and lack comprehensive evaluation or validation. Here we use a CRISPR/cas9 knockin-based saturation genome editing (SGE) MAVE to evaluate the functional consequences of all SNVs in the 12 exons of *BRCA2* encoding the BRCA2 DBD, which is the sole location of known pathogenic missense variants in this gene. The results were combined with other sources of genetic and clinical evidence in an ACMG/AMP model for classification of variants as pathogenic or benign and for development of a comprehensive reference for clinical management of individuals found to carry any VUS in this domain.

## Results

### Saturation genome editing (SGE) of *BRCA2*

SGE of exons 15 to 26 of *BRCA2* (MANE transcript ENST00000380152.8), encoding the BRCA2 DBD (amino acids 2481-3186) hotspot region for pathogenic missense variants, was performed in the haploid human HAP1 cell line in order to insert all possible DBD SNVs into the endogenous BRCA2 gene and to assess the functional impact of all variants on cell viability. *BRCA2* has been identified as an essential gene in HAP1 cells^20,21^.

On this basis, the influence of BRCA2 deficiency on viability of these cells, sorted for haploidy, was confirmed by luminescence-based ATPase assay analysis of cells at different time points after CRISPR/Cas9 targeting of exon 19 with two guide RNAs (sgRNAs) (**Extended Data** Fig. 1). Next, individual coding exons along with 10bp of adjacent intronic nucleotides, except for exons 18 and 25, which were divided into two separate regions, were selected as SGE target regions in the MAVE experiments (**Fig. 1A**). The most efficient sgRNA for each target region was selected and cloned into a Cas9 expressing plasmid (**Fig. 1B, Supplementary Table 1, 2**). Site-saturation mutagenesis (SSM) libraries for each of the 14 target regions were designed to target every nucleotide and were generated by site directed mutagenesis using NNN-tailed PCR primers (**Fig. 1B, Supplementary Table 2**). The combined target libraries contained 7233 of 7239 (99.9%) possible SNVs, with six missense variants missing from exon 16. For each target region 2 million HAP1 cells were cultured, cotransfected on Day (D) 0 with sgRNA/Cas9 and variant library plasmids, selected with puromycin for three days and sampled on D5 (24 hrs after puromycin selection) and D14. Genomic DNA (gDNA) from each timepoint was harvested and subjected to amplicon-based massively parallel sequencing (**Fig.1B**). Paired end sequencing reads were used to estimate individual SNV counts at each timepoint. The average sequencing depth for each variant was 3694 reads for D5 replicates and 3046 reads for D14.

**Fig. 1.**
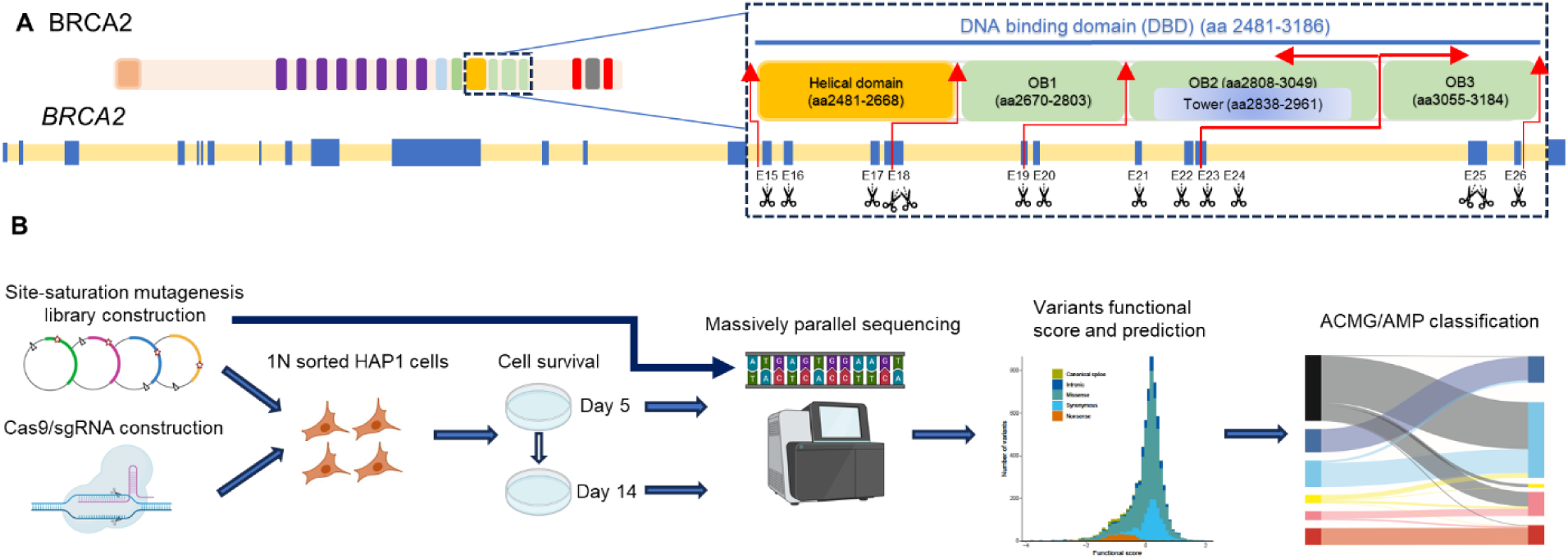
Schematic overview of a saturation genome editing (SGE) multiplexed assay of variant effect (MAVE) of all single nucleotide variants (SNVs) in the BRCA2 DNA Binding Domain (DBD). **A**: Design of the SGE experiment target regions. All possible SNVs were introduced and assessed in exons 15-26 encoding the BRCA2 DBD Helical (AA 2481-2667), OB1 (AA 2670-2803), OB2 (AA 2808-3049, including Tower subdomain (AA 2838-2961)), and OB3 (AA 3055-3184) domains, along with 10bp of adjacent intronic nucleotides for each exon. Exons 18 and 25 were divided into two regions, resulting in 14 total target regions. **B**: Schematic illustrating the SGE workflow. In each target region, respective SNV library was generated to saturate the region with all possible SNVs. The SNV library was transfected with corresponding Cas9/gRNA construct into HAP1 haploid cells. Genomic DNA (gDNA) was extracted at post-transfection day 5 and 14, targeted region was amplified and barcoded for targeted gDNA sequencing. SNV abundance was evaluated and normalized to generate functional scores for all SNVs. An ACMG/AMP classification model was applied to formally classify SNVs based on the MAVE functional result and other evidence.

### Functional analysis of variant effects

Individual SNVs were labeled using BioR and the Ensembl Variant Effect Predictor (VEP). Log2 Fold Changes (LFCs) of D14 to D5 ratios were calculated for each independent experiment. No position dependent effects within target regions were noted. SNV LFCs for each replicate were normalized within each exon and across all exons relative to median synonymous and median nonsense SNV scores. Replicates of individual SNVs with LFCs>1 relative to mean replicate values were excluded and LFCs were recalculated (**Fig. 2A, Extended Data** Fig. 2**, Supplementary Table 3**). Functional scores for 7013 (96.9%) SNVs were obtained that encoded 339 nonsense, 1377 synonymous, and 4594 missense, along with 142 canonical splice site (+/1-2) and 561 intronic variants (+/-3 to +/-10), (**Fig. 2A, Extended Data Table 1**).

**Fig. 2.**
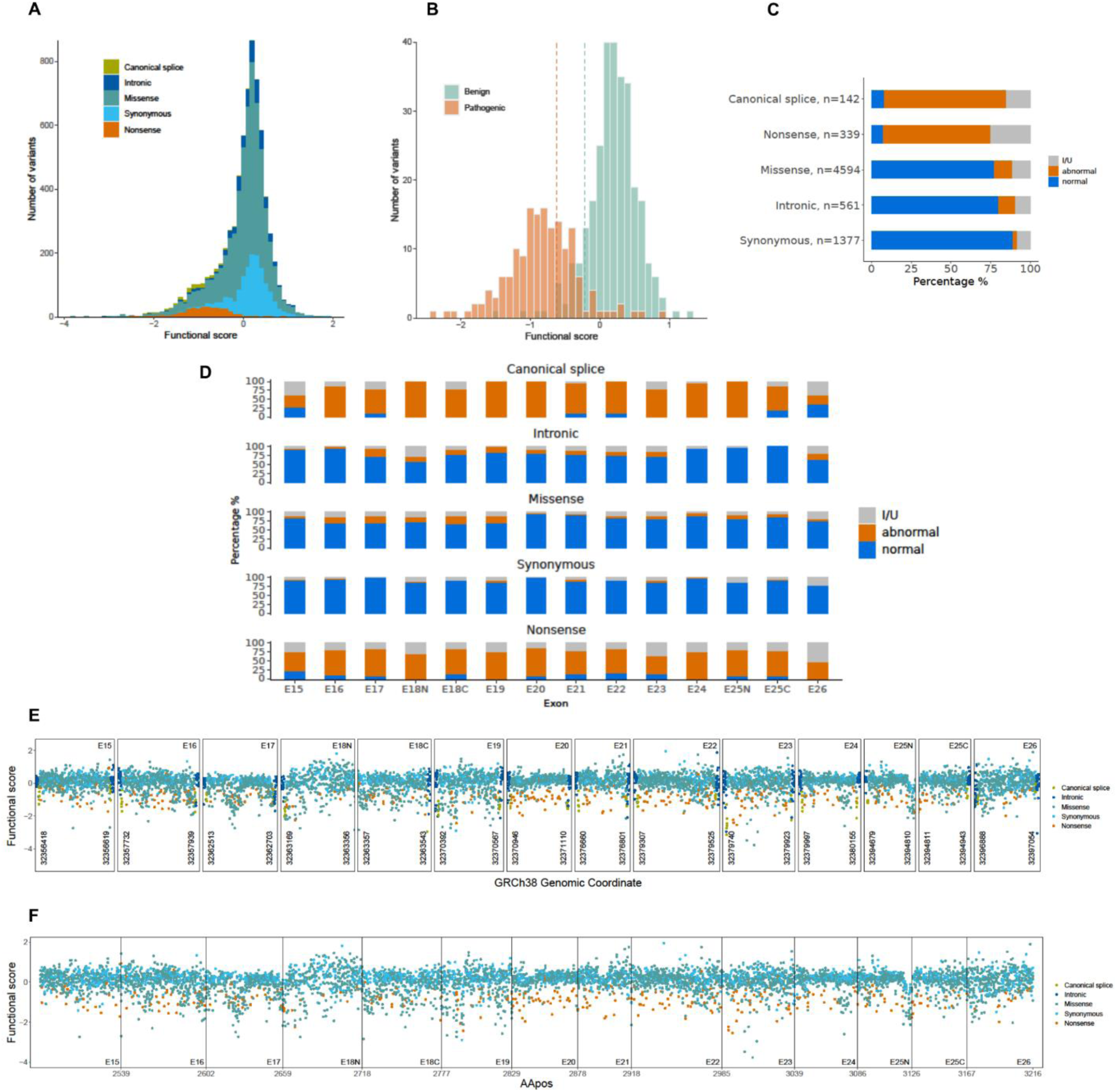
Functional annotation of BRCA2 SNVs. **A**: Distribution of functional scores of 7013 SNVs evaluated in BRCA2, colored by variant types. **B**: Functional score thresholds were determined by 95% prevalence of functional scores from curated ClinVar established pathogenic (n=162) and benign (n=259) non-splicing variants. Functionally abnormal variants scored ≤-0.615. Functionally normal variants scored ≥-0.225. **C**: Bar chart illustrating the number of SNVs within each variant category. Color indicates functional category (I/U, intermediate/uncertain). **D**: Functional categorization of variant types in each exon. **E**: Functional score map of SNVs by nucleotide position in 14 target regions, including exons and flanking canonical splice site and intronic variants, colored by type of variant effect. **F**: Functional score map of SNVs from 12 exons by encoded amino acid position, colored by type of variant effect. Exons 18 and 25 were both divided into two regions (N-terminus and C-terminus).

**Table 1:**
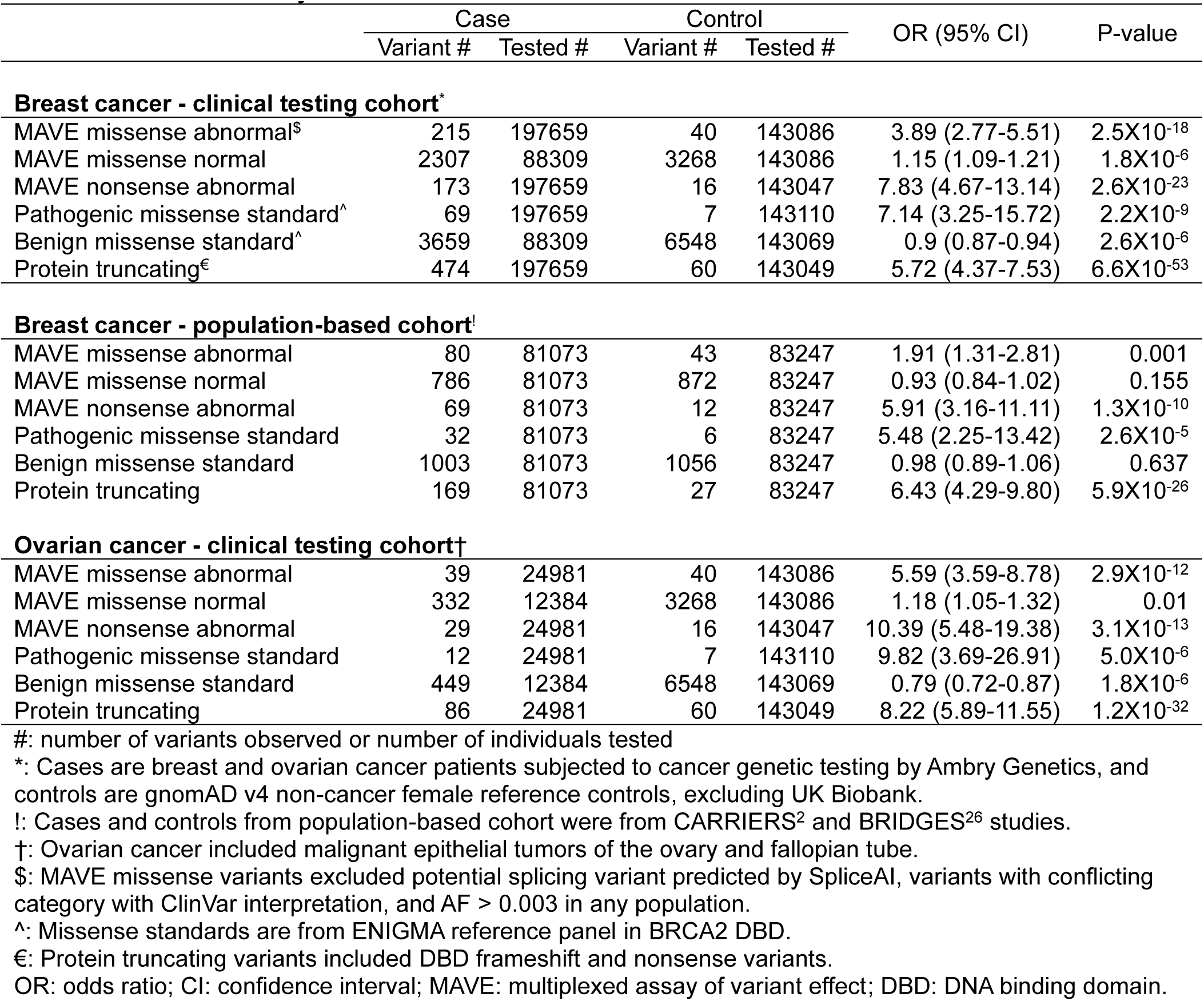
Association analysis of MAVE characterized variants with breast and ovarian cancer.

|  | Case |  | Control |  | OR (95% CI) | P-value |
| --- | --- | --- | --- | --- | --- | --- |
|  | Variant # | Tested # | Variant # | Tested # |  |  |
| **Breast cancer - clinical testing cohort\*** |  |  |  |  |  |  |
| MAVE missense abnormal<sup>§</sup> | 215 | 197659 | 40 | 143086 | 3.89 (2.77-5.51) | 2.5X10<sup>-18</sup> |
| MAVE missense normal | 2307 | 88309 | 3268 | 143086 | 1.15 (1.09-1.21) | 1.8X10<sup>-6</sup> |
| MAVE nonsense abnormal | 173 | 197659 | 16 | 143047 | 7.83 (4.67-13.14) | 2.6X10<sup>-23</sup> |
| Pathogenic missense standard<sup>^</sup> | 69 | 197659 | 7 | 143110 | 7.14 (3.25-15.72) | 2.2X10<sup>-9</sup> |
| Benign missense standard<sup>^</sup> | 3659 | 88309 | 6548 | 143069 | 0.9 (0.87-0.94) | 2.6X10<sup>-6</sup> |
| Protein truncating<sup>€</sup> | 474 | 197659 | 60 | 143049 | 5.72 (4.37-7.53) | 6.6X10<sup>-53</sup> |
| **Breast cancer - population-based cohort<sup>!</sup>** |  |  |  |  |  |  |
| MAVE missense abnormal | 80 | 81073 | 43 | 83247 | 1.91 (1.31-2.81) | 0.001 |
| MAVE missense normal | 786 | 81073 | 872 | 83247 | 0.93 (0.84-1.02) | 0.155 |
| MAVE nonsense abnormal | 69 | 81073 | 12 | 83247 | 5.91 (3.16-11.11) | 1.3X10<sup>-10</sup> |
| Pathogenic missense standard | 32 | 81073 | 6 | 83247 | 5.48 (2.25-13.42) | 2.6X10<sup>-5</sup> |
| Benign missense standard | 1003 | 81073 | 1056 | 83247 | 0.98 (0.89-1.06) | 0.637 |
| Protein truncating | 169 | 81073 | 27 | 83247 | 6.43 (4.29-9.80) | 5.9X10<sup>-26</sup> |
| **Ovarian cancer - clinical testing cohort†** |  |  |  |  |  |  |
| MAVE missense abnormal | 39 | 24981 | 40 | 143086 | 5.59 (3.59-8.78) | 2.9X10<sup>-12</sup> |
| MAVE missense normal | 332 | 12384 | 3268 | 143086 | 1.18 (1.05-1.32) | 0.01 |
| MAVE nonsense abnormal | 29 | 24981 | 16 | 143047 | 10.39 (5.48-19.38) | 3.1X10<sup>-13</sup> |
| Pathogenic missense standard | 12 | 24981 | 7 | 143110 | 9.82 (3.69-26.91) | 5.0X10<sup>-6</sup> |
| Benign missense standard | 449 | 12384 | 6548 | 143069 | 0.79 (0.72-0.87) | 1.8X10<sup>-6</sup> |
| Protein truncating | 86 | 24981 | 60 | 143049 | 8.22 (5.89-11.55) | 1.2X10<sup>-32</sup> |

#: number of variants observed or number of individuals tested
\*: Cases are breast and ovarian cancer patients subjected to cancer genetic testing by Ambry Genetics, and controls are gnomAD v4 non-cancer female reference controls, excluding UK Biobank.

To standardize the assay relative to variants of known effect and to establish functional score thresholds for functionally abnormal and functionally normal SNVs, 162 pathogenic and 259 benign variants with no effects on splicing and reported in ClinVar with consistent findings from at least two ClinGen-approved testing laboratories or from the ENIGMA (Evidence-based Network for the Interpretation of Germline Mutant Alleles) *BRCA1/2* expert panel were used. Based on 95% prevalence, a functionally abnormal functional score threshold corresponding to ClinVar reported pathogenicity was set at <= –0.615 and a benign and functionally normal threshold was set at >= –0.225 for all SNVs in exons 15-22, 24 and 25 (**Fig. 2B**). SNVs in exons 23 and 26 were excluded from this model because of relatively poor performance (<90% sensitivity and specificity) relative to ClinVar known pathogenic and benign standards (**Fig. 2B**). In order to incorporate variants from exons 23 and 26 in subsequent analyses of the MAVE dataset, a secondary analysis of all exons (15 to 26) was performed that yielded 95% prevalence thresholds of –0.845 for abnormal SNVs and –0.260 for normal SNVs. In this secondary model, 47.5% of nonsense variants in exons 23 and 26 were categorized as Intermediate/uncertain compared to only 22.7% in the high quality (exons 15-22, 24 and 25) model. Results for all SNVs in exons 23 and 26 in this more conservative model were subsequently extracted and combined with results from the initial (exons 15-22, 24 and 25) model to yield a primary dataset for the MAVE study (**Extended Data Table 2, Supplementary Table 3**).

Validation of the MAVE thresholds and categorization of SNVs was performed by comparison with nonsense and synonymous variants. Only 19 of 260 (7.3%) nonsense variants were scored as functionally normal by MAVE, whereas 22 of 1134 (1.9%) synonymous variants were scored as functionally abnormal (**Extended Data Table 3**). Furthermore, only 6 of 106 (5.7%) functionally abnormal missense variants from previous HDR assays^11–13,22^ were scored as functionally normal in the MAVE assay, whereas 11 of 263 (4.2%) of HDR functionally normal were scored as functionally abnormal by MAVE (**Extended Data Table 3**). Finally, none of 16 missense variants encoded by SNVs that were classified as pathogenic missense standards by the ENIGMA consortium and ClinGen VCEP for *BRCA2* variant classification were scored as functionally normal, and only 2 of 50 (4%) ENIGMA classified benign missense variants were scored as functionally abnormal by MAVE (**Extended Data Table 3**). Thus, error rates for the MAVE functional assays, as defined by three independent validation datasets, were very low.

Based on these MAVE thresholds, all 7013 SNVs were categorized as functionally abnormal, functionally normal or intermediate/uncertain (I/U) (**Fig. 2C, Extended Table 2**). In total, 522 SNVs encoding missense, 228 nonsense, 109 canonical splice, 60 intronic, and 36 synonymous variants were scored as functionally abnormal (**Fig. 2C, Extended Data Table 2**). In contrast, 3524 SNVs encoding missense, 24 nonsense, 11 canonical splice, 446 intronic, and 1219 synonymous variants were scored as functionally normal. The remaining 834 variants remained undefined in the I/U category (**Fig. 2C, Extended Data Table 2**). Functionally abnormal missense SNVs were enriched in exons 16-19, with abnormal rates ranging from 17-22%, whereas rates were as low as 2% in exons 20 and 21 that encode the OB2 and Tower domains (**Fig. 2D, Extended Data Table 2**). Missense SNV I/U rates ranged from 5-21% across the region. Among 142 variants in +1/2 and –1/2 canonical splice site positions, only 11 yield normal function including six with modifications in the leaky +2 position (**Fig. 2C, E, Extended Data Table 2**). Furthermore, terminal G nucleotides that contribute to splice site activity were present in most exons. Of 27 SNVs in these terminal G alleles, 18 had abnormal function and only 4 had normal function (**Fig. 2F, Extended Data Table 2**). In addition, 60 intronic SNVs had abnormal activity and another 55 were I/U (**Fig. 2C, Extended Data Table 2, Supplementary Table 3**). Thus, even though RNA studies were not performed, the MAVE study may have identified a large number of variants that influence RNA splicing.

### Correlation with DBD architecture

To gain insight into the mechanisms by which the functionally abnormal SNVs disrupt BRCA2 activity and to evaluate whether variant hotspots are present in the BRCA2 DBD, the influence of the functionally abnormal SNVs on protein structure was assessed. MAVE data from all exons, including exons 23 and 26, was used. Functionally abnormal missense variants were enriched in the exons encoding the Helical and OB1 domains (16.7%, 322 of 1926 missense SNVs) but were less common (11.1%, 173 of 1561 missense SNVs) in the OB2 and OB3 domains (p=2.1×10^-6^), and very infrequent (1.9%, 15 of 803 missense SNVs) in the Tower protein domain (**Extended Data Table 4**). In addition, among the 522 of 4594 (11.4%) missense SNVs in the DBD that were functionally abnormal, 186 (35.6%) were in the Helical domain, 136 (26.1%) in OB1, 85 (16.3%) in OB2, 15 (2.9%) in the Tower, 93 (17.8%) in OB3 and another 7 in linker regions (**Fig. 3A, Extended Data Table 4**). These findings were consistent with HDR assay results for 462 DBD missense variants which showed an enrichment for functionally abnormal missense alterations in the Helical, OB1 and OB3 domains^13^. Approximately 75% of the MAVE functionally abnormal SNVs were located in α-helices and β-sheet structures needed to maintain essential 3D folding. Few (n=128) functionally abnormal missense variants were located in disordered or loop regions. Functionally abnormal missense alterations were observed in 52% (26 of 50) of residues known to interact with the DSS1 stabilizing protein and 12% (2 of 17) of residues involved in single strand DNA interactions suggesting a strong requirement for these interactions for BRCA2 activity. It was also noted that 288 of 503 (57%) functionally abnormal missense alterations resulted in charge changes or loss or gain of Proline residues (**Supplementary Table 3**).

**Fig. 3.**
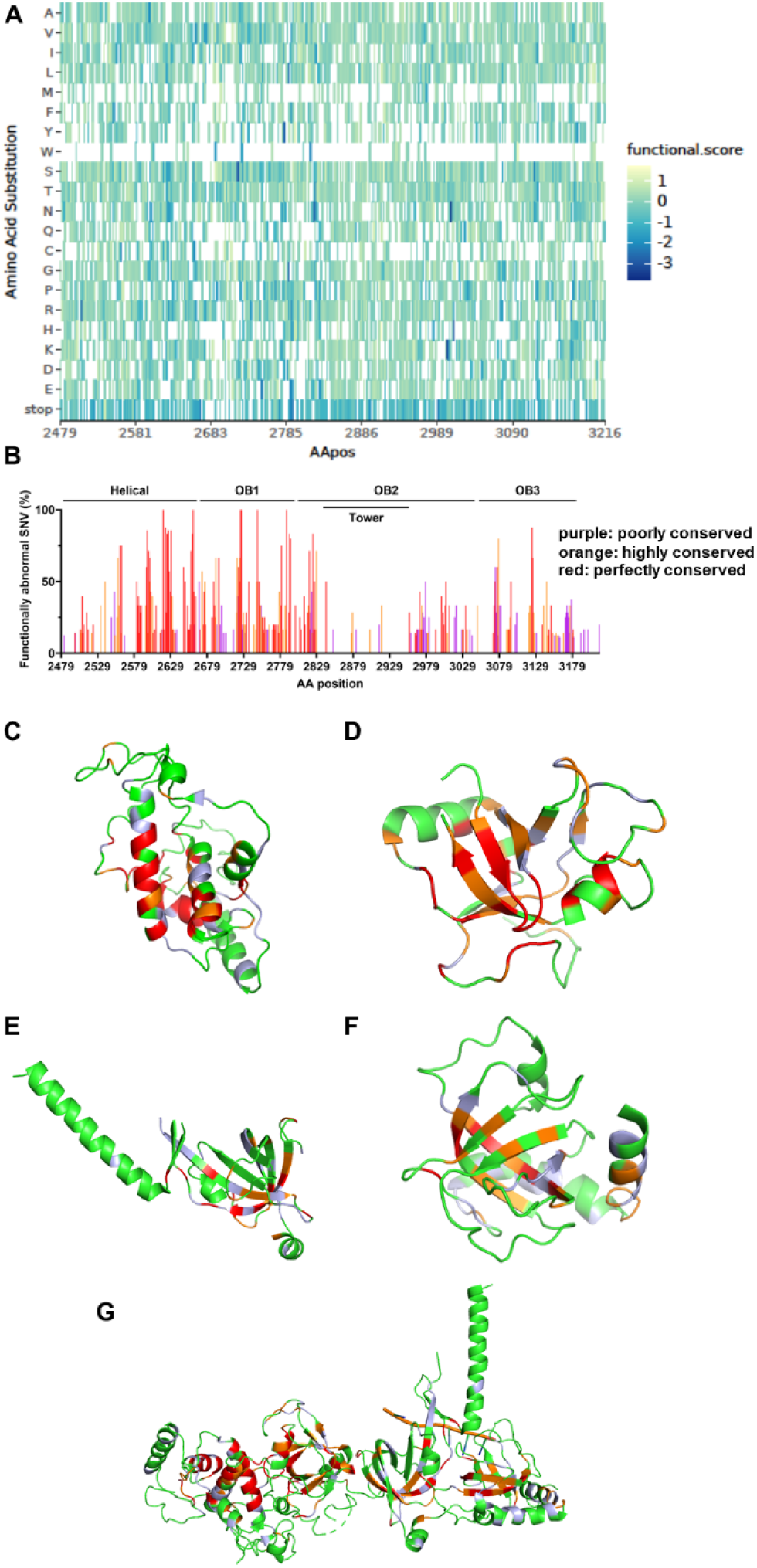
Functional effect of SNVs on BRCA2 protein. **A**: Heatmap of average functional scores for each amino acid substitution encoded by SNVs. **B**: Cross-species sequence conservation from pufferfish to homo sapiens (n=10) relative to frequency of functionally abnormal SNVs accounting for each missense alteration (perfectly conserved: 100% identity across 10 species; highly conserved: 80% or 90% identity across 10 species; poorly conserved: ≤70% identity across species). **C, D, E, F, G**: BRCA2 3-dimensional protein ribbon diagram showing missense alterations, encoded by functionally abnormal SNVs, in the Helical (**C**), OB1 (**D**), OB2 (**E**), OB3 (**F**) domains, and the BRCA2-DSS1-ssDNA complex (PBD 1MJE) (**G**). Color denotes the frequency of functionally abnormal SNVs accounting for each missense alteration (Green: 0%; purple: <20%; orange: 20-39.9%; red: ≥40%. The subdomains were oriented to maximize views of the functionally abnormal missense alterations. The BRCA2-DSS1-ssDNA complex is shown from N-terminus (left) to C-terminus (right). ssDNA: single-stranded DNA.

The BRCA2 DBD is highly conserved from pufferfish to *Homo sapiens*, with many residues perfectly conserved across species. Among 213 of 220 perfectly conserved residues evaluated, at least one functionally abnormal SNV resulting in a missense substitution was observed in 121 (57%) (**Fig. 3B, Extended Data Table 4**). Another 24% of these conserved residues were influenced by Intermediate/uncertain variants, which may have partial effects on BRCA2 function (**Extended Data Table 4**). Consistent with the 3D structural model, the majority of the functionally abnormal variants in conserved residues were located in the Helical (50 of 121, 41.3%) and OB1 (32 of 121, 26.4%) domains (**Fig. 3B-G**). Only 13 of 51 amino acids in OB3 with at least one functionally abnormal missense change were from the 19 fully conserved residues in this domain (**Extended Data Table 4**).

### Comparisons with functional predictors

Several methods that have been used to assess the influence of *BRCA2* missense variants on protein function have been selected by the ENIGMA consortium and the ClinGen VCEP as appropriate for utilization in variant clinical classification models. As noted above, a strong correlation was found between the MAVE data and results for 462 BRCA2 DBD missense variants evaluated using a cell-based HDR assay. Importantly MAVE data distinguished between HDR abnormal, normal and hypomorph/intermediate categories (p<0.05) (**Fig. 4A**).

**Fig. 4.**
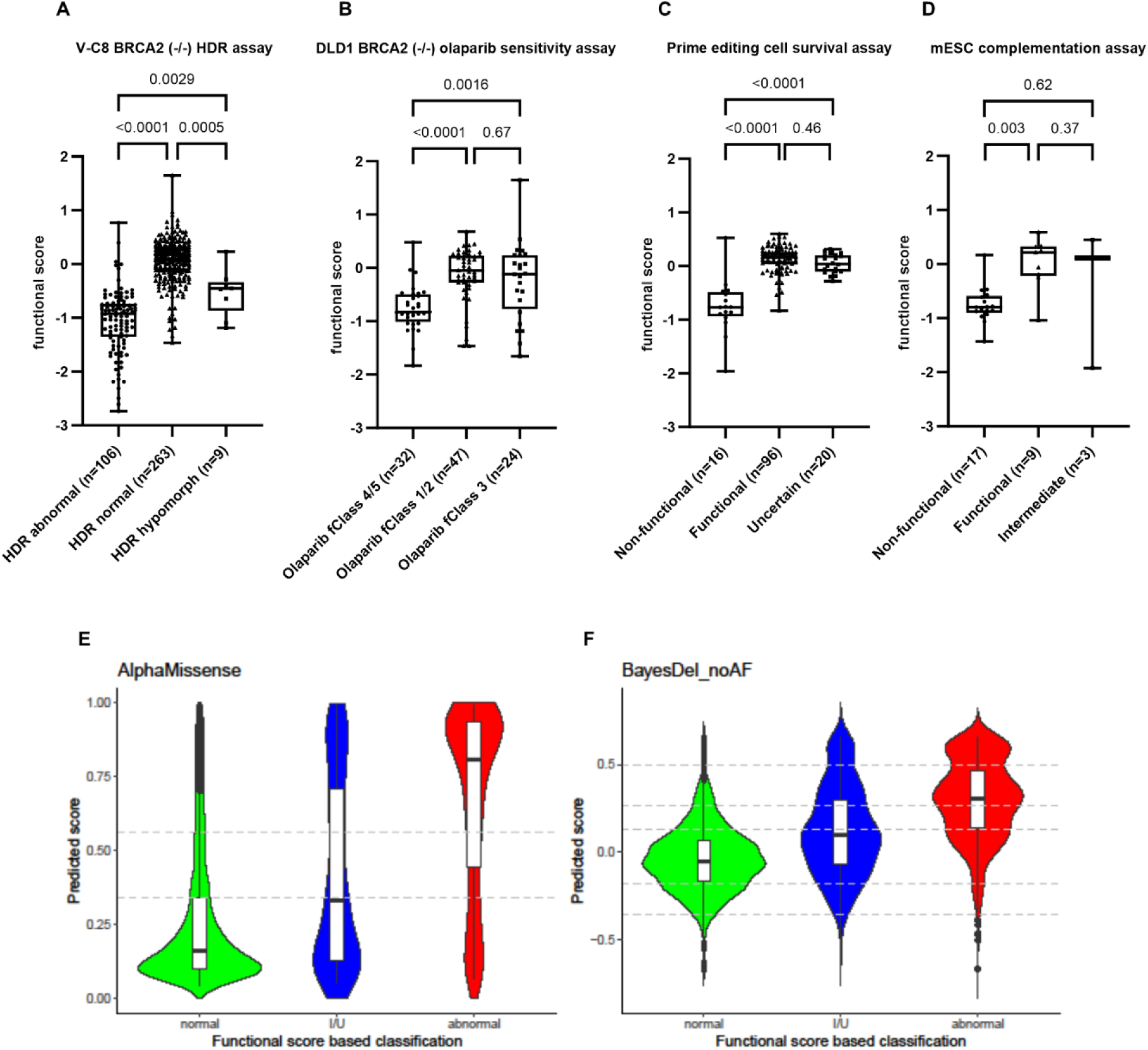
Comparison of BRCA2 MAVE with functional assays and *in silico* predictors. Functional scores of SNVs encoding missense compared with: **A**: BRCA2 (-/-) V-C8 homology directed repair (HDR) assay; **B**: DLD1 BRCA2 (-/-) olaparib sensitivity assay; **C**: prime editing-based haploid cell survival assay; **D**: murine *brca2-/-* embryonic stem cell (mESC) complementation assay. Numbers of variant of each type resulting from the individual assays are shown. One-way ANOVA tests were used to calculate statistical significance of differences between functional categories. **E**: AlphaMissense in silico prediction scores compared to MAVE functional scores of SNVs resulting in missense alterations (n=3940) (likely pathogenic >0.564; likely benign <0.34; ambiguous: <=0.564, >=0.34). Differences between functionally abnormal and normal variants were statistically significant (Wilcoxon rank-sum test, p=4.5×10^-122^). **F**: BayesDel in silico prediction scores compared to MAVE functional scores of SNVs resulting in missense alterations. Differences between functionally abnormal and normal variants were statistically significant (Wilcoxon rank-sum test, p=1.3×10^-220^). BayesDel thresholds were defined according to ACMG/AMP classification rules evidence strength: PP3 (pathogenic): Strong: ≥0.5; Moderate: 0.27 to 0.5; Supporting: 0.13 to 0.27. BP4 (benign): Supporting: –0.36 to –0.18; Moderate: ≤-0.36.

Similarly, MAVE data effectively distinguished between Class 4,5 (functionally abnormal) and Class 3 (uncertain) (p=0.0016) and between Class 4,5 and Class 1,2 (functionally normal) (p<0.0001) in an olaparib PARPi response assay (**Fig. 4B**)^23^. Both of these assays involved overexpression of BRCA2 cDNAs in BRCA2 deficient cells. However, the MAVE data also discriminated effectively between non-functional and functional (p<0.0001) or uncertain (p<0.0001) variants in an endogenously targeted prime editing study limited to exons 15 and 17 (**Fig. 4C**)^17^. Finally, the MAVE data successfully distinguished between non-functional and functional (p=0.003) missense variants in a small embryonic stem cell complementation assay^24^. Thus, the MAVE analysis is highly consistent with results from several other small scale functional assays (**Fig. 4A-D, Supplementary Table 4**), but improves upon these assays by identifying Class 3/uncertain/intermediate variants from these assays as predominantly functionally normal.

Next, comparisons between the MAVE results and *in silico* prediction methods were performed. BayesDel is the predictor currently used by the ClinGen *BRCA1/2* VCEP for curation of *BRCA1/2* variants, whereas the AlphaMissense method is based on unsupervised protein language modeling and incorporates structural context from an AlphaFold-derived system ^22^ (**Fig. 4E-F**, **Supplementary Table 5**). The sensitivity of BayesDel for functionally abnormal SNVs was 0.95, but the specificity for functionally normal SNVs was 0.58, suggesting that many false positives can be predicted by the BayesDel model. In contract, AlphaMissense yielded 0.69 sensitivity and 0.75 specificity (**Supplementary Table 5**). Both models are effective predictors of MAVE assay results. The utility of these prediction models in the presence of MAVE data for 97% of all SNVs in the BRCA2 DBD must now be questioned.

### Cancer risks associated with functionally abnormal variants

To understand the contributions to cancer risk of the functionally abnormal SNVs encoding missense variants associations between pooled functionally abnormal variants and breast and ovarian cancer were evaluated. Specifically, the frequency of the pooled functionally abnormal missense variants in female, primary breast cancer cases subjected to hereditary cancer genetic testing by Ambry Genetics from 2012-2021 was compared to the frequency in female, non-cancer reference controls from gnomAD V4, excluding UK Biobank^25^. The functionally abnormal missense SNVs yielded an odds ratio (OR) for breast cancer of 3.89 (95%CI: 2.77-5.51) (**Table 1**). This odds ratio was attenuated compared to results for MAVE nonsense variants (OR 7.83, 95%CI: 4.67-13.14), established pathogenic missense variants (OR 7.14, 95%CI: 3.25-15.72), and DBD protein truncating variants (OR 5.72, 95%CI: 4.37-7.53), although the differences were not significantly different (**Table 1**). Importantly, the missense SNVs scored as functionally normal in the MAVE study were not associated with clinically relevant (OR>2) increased breast cancer risk (OR 1.15, 95%CI: 1.09-1.21) and were similar to established benign variants (OR 0.9, 95%CI: 0.87-0.94) (**Table 1**). Furthermore, functionally abnormal missense SNVs yielded similar associations in Non-Finnish Europeans (NFE) (OR 4.99, 95%CI: 3.05-8.29) and African Americans (AA) (OR 4.67, 95%CI: 1.73-13.08), whereas functionally normal missense variants were not associated with elevated breast cancer risk in NFE (OR 1.07, 95%CI: 1.00-1.15) or in AA (OR 1.03, 95%CI: 0.9-1.18) (**Extended Data Table 5**).

Additional analyses using case-control data from the CARRIERS and BRIDGES population-based breast cancer studies^2,26^ yielded consistent findings, although the ORs were attenuated due to the population-based nature of the cases and controls (**Table 1**). Functionally abnormal missense SNVs again showed attenuated risks relative to nonsense and known pathogenic missense variants (**Table 1**). A parallel analysis of associations with ovarian cancer risk found that the pooled functionally abnormal missense SNVs were associated with substantially increased risks of ovarian cancer (OR 5.59, 95%CI: 3.59-8.78) and as with breast cancer, the missense SNVs showed attenuated risks relative to nonsense variants (**Table 1).**

### Clinical classification of MAVE SNVs

As noted above, functional data for SNVs must be integrated into classification models in order to determine the clinical relevance of each variant and to use the added information for clinical management of variant carriers. An ACMG/AMP classification framework for *BRCA2* variant classification that incorporated functional HDR data and used point scoring rather than qualitative estimates for each classification rule was previously applied to 442 variants^13^. Here, a similar ClinGen/ACMG/AMP classification framework was applied to the MAVE SNVs (**Supplementary Table 6, Fig. 5**). Initially, all nonsense and canonical splice site variants were scored as PVS1 and directly attributed 8 points for a pathogenic classification^27^. Next, the odds of pathogenicity for the MAVE functional assay were assessed based on the ClinVar positive and negative controls. The functionally abnormal threshold yielded odds of pathogenicity of 43:1 for abnormal variants, equating to “strong” evidence of pathogenicity for the PS3 functional assay ACMG/AMP rule^28^. Conversely the functionally normal threshold yielded odds of benign of 0.09:1 for normal variants and “moderate” evidence of benign for the BS3 functional assay rule.

**Fig. 5.**
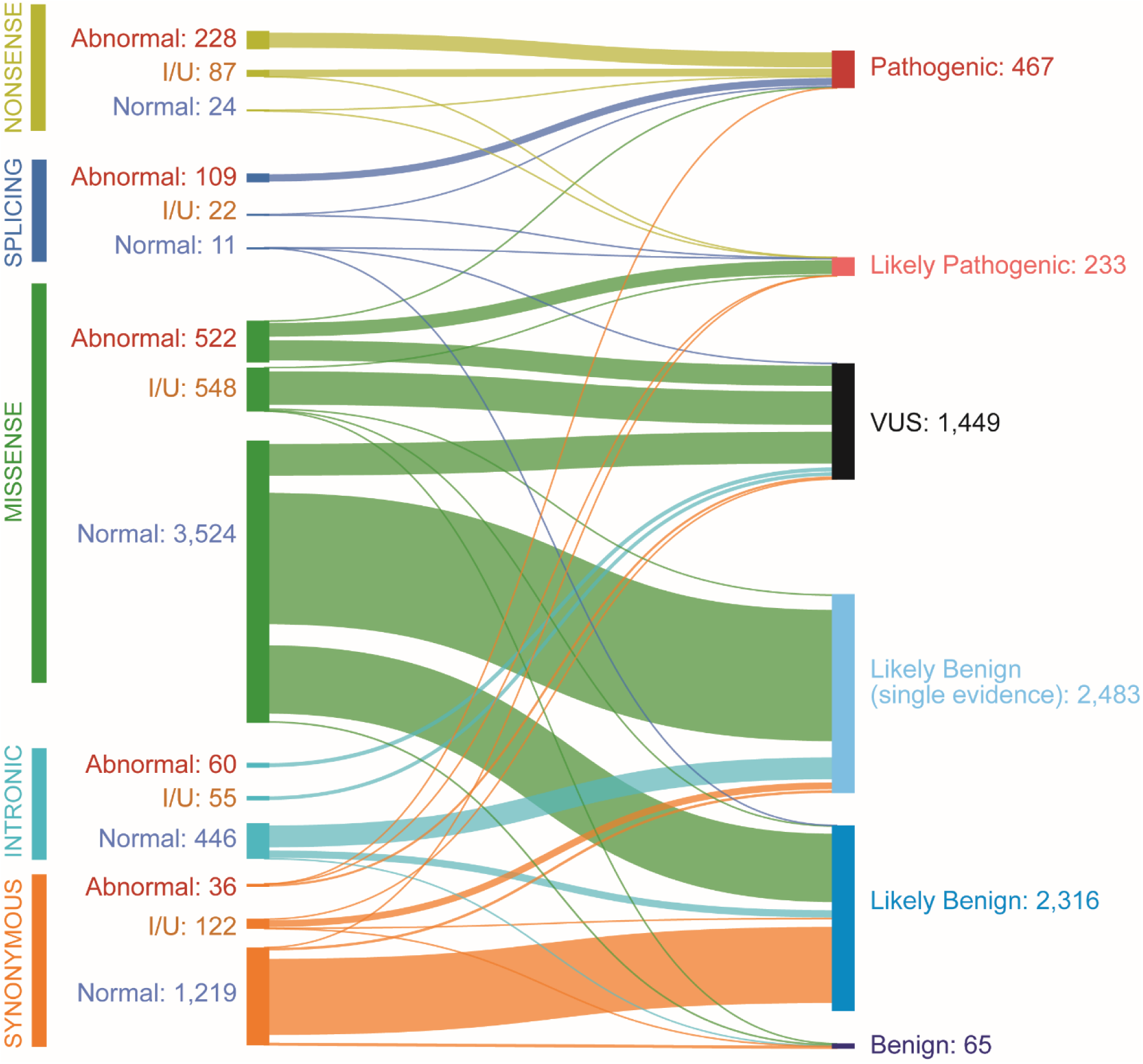
Clinical classification of MAVE SNVs. Sankey plot illustrating the clinical classification of SNVs after applying a ClinGen/ACMG/AMP framework. Of 7013 MAVE evaluated SNVs, 467 were classified as pathogenic, 233 as likely pathogenic, 65 as benign, 4799 as likely benign, and 1449 remained as VUS. VUS: variants of uncertain significance.

This rule was subsequently applied to all SNVs and +4 points for PS3 pathogenicity were awarded to all functionally abnormal variants, whereas –2 points for BS3 in favor of benignity were awarded to functionally normal variants. I/U variants received 0 points. PS3 was combined with PP3 BayesDel *in silico* predictions, PS4 case-control associations, PM3 *in trans* – Fanconi Anemia for evidence in favor of pathogenicity (**Supplementary Table 6**). In contrast, BS3 normal functional evidence was combined with BA1 and BS1 population frequencies, BP4 *in silico* prediction for benign, and BP2 *in trans*-healthy for evidence in favor of benign effects (**Supplementary Table 6**). PVS1 evidence for pathogenicity (+8 points) was applied to nonsense and canonical splice site SNVs, except for five variants at +2 canonical splice sites with potentially leaky splicing effects **Supplementary Table 6**).

The integration of PVS1 with other evidence codes in the ACMG/AMP model scored 7 nonsense and 13 canonical splice site SNVs as likely pathogenic (LP), whereas 332 nonsense and 124 canonical splice SNVs were classified as pathogenic (P). Among the 4594 missense SNVs evaluated in the MAVE study, 220 were LP/P, 3084 were likely benign/benign (LB/B) and 1290 remained as VUS (**Fig. 5, Extended Data Table 6**). For all SNVs in the MAVE study, 4864 were classified as B/LB, 700 as P/LP and 1449 as VUS. To evaluate the impact of the MAVE study on variant classification, the MAVE results were combined with classifications from ClinVar and ENIGMA. Of 5043 SNVs classified as B/LB, ClinVar/ENIGMA accounted for 993 (19.7%) and MAVE uniquely accounted for 4050 80.3%; among 775 classified as P/LP, ClinVar/ENIGMA accounted for 417 (53.8%) and MAVE uniquely accounted for 358 (46.2%) (**Fig. 5, Extended Data Table 6, Supplementary Table 6**). In addition, of 320 SNVs with discordant classifications in ClinVar (P/LP vs VUS, or B/LB vs VUS), 230 (71.9%) were resolved by MAVE-based classification were resolved as B/LB or P/LP, while 50 discordances remained as VUS. Thus, the MAVE study had a tremendous impact on classification of variants in *BRCA2* exons 15-26 that encode the DBD.

### Phenotypic characteristics associated with functionally abnormal SNVs

The average age of breast cancer diagnosis among individuals from the clinical testing cohort with either functionally abnormal missense SNVs or nonsense SNVs was 52 years, consistent with an earlier age at diagnosis due to predisposing alleles in BRCA2. A significant difference in family history of breast cancer, defined as any first or second degree relative with disease, was also observed for individuals carrying germline functionally abnormal missense SNVs (126 of 161 (78.3%)) relative to functionally normal SNVs (1644 of 2397 (68.6%)) (p=0.01). Lifetime risk estimates for breast cancer were estimated using ORs from the current study and SEER rates of disease. The abnormal missense SNVs had an estimated lifetime risk of 46% to age 80, similar to the 55% risks for DBD nonsense and protein truncating variants, and substantially greater than the 12% risks for normal missense SNVs. Analyses were restricted to non-Hispanic whites because of limited statistical power for estimation of breast cancer ORs for other populations.

Loss of heterozygosity (LOH) of BRCA2 in tumors was evaluated to assess whether the functionally abnormal SNVs in the MAVE assay may be drivers of tumor development. LOH at *BRCA2* was evaluated in 50,000 breast, ovarian, prostate and pancreatic tumor samples with greater than 40% tumor content that had been sequenced using a cancer gene panel in the IMPACT study^29^. Among tumors with *BRCA2* abnormal SNVs of any type, 18 of 21 (86%) displayed LOH. Similarly, LOH was found in 12 of 15 (80%) tumors associated with abnormal missense and nonsense SNVs. In contrast, 54 of 209 (26%) tumors with functionally normal variants displayed LOH (p=9.4×10^-8^). Thus, functionally abnormal variants appeared to be selected for loss of the wildtype BRCA2 allele and inactivation of BRCA2, consistent with a role for the BRCA2 abnormal SNVs as drivers of tumor initiation.

## Discussion

The functional evaluation of variants in BRCA2 has been an active area of research because of the high risks of several cancers (breast, ovarian, prostate, pancreatic, endometrial and cholangiocarcinoma) associated with inactivating variants in BRCA2, the large number of VUS in BRCA2 that may only be classified upon the inclusion of functional evidence, and the insight into BRCA2 function and biology that can be gained from such studies. However, to date only 557 missense variants in the BRCA2 DBD have been evaluated by well-established functional assays^7,11–13,23,24^, and only 418 of these have been clinically classified as P/LP or B/LB. The substantial number of identified variants with clinical uncertainty has necessitated more rapid functional characterization. Herein, an SGE study in human haploid cells was used to functionally evaluate the effects of all BRCA2 SNVs in the BRCA2 DBD pathogenic missense variant hotspot on BRCA2 activity, as measured by cell viability. Functional scores were obtained for 7013 SNVs (97% of all possible SNVs) from 12 coding exons and flanking intronic sequences. Using ClinVar established clinically pathogenic and benign variants as standards, 95% thresholds for functionally abnormal and normal categories were defined and 955 SNVs were identified as abnormal, 834 as I/U and 5224 as normal. While more than 600 DBD SNVs have previously been evaluated by other functional assays, the current study is the first to establish a sequence-function map for nearly all possible SNVs in the BRCA2 DBD. The functional results from this study provided structural insight into the BRCA2 protein. Functionally abnormal missense SNVs were spread across the Helical, OB1, OB2 and OB3 domains, with a slight enrichment in the Helical and OB1 subdomains (60%, 322 of 522). Interestingly, 15 abnormal missense SNVs were found in the Tower domain where no established pathogenic missense variants have been reported. This finding confirms that the Tower domain is required for normal BRCA2 function, establishes that the Tower is not a cold spot for disease causing variants, and suggests that missense variants in the domain are associated with increased risk of cancer. Likewise the enrichment of abnormal missense SNV in DSS1 binding sites but not in single strand DNA binding sites suggests that the stability of this domain rather than interactions with DNA is most important for BRCA2 function.

Importantly, the MAVE functional data do not in themselves classify the clinical relevance of any variants. This can only be achieved by incorporating the functional data into classification models. Thus, after establishing that the MAVE assay conferred “strong evidence” of pathogenicity under the PS3/BS3 rule the evidence was integrated with other genetic and clinical data into an ACMG/AMP/ClinVar classification model for *BRCA2* variants. The outcome was that 220 missense SNVs were classified as P or LP and 3084 were classified as B or LB. The remaining 1290 SNVs remained as VUS. However, with the availability of the MAVE data, it seems likely that many of these variants will be classified as P/LP or B/LB in the future following the addition of data from other sources.

While 1410 *BRCA2* DBD SNVs had previously been classified by ClinVar as P/LP (n=417) or B/LB (n=993), the MAVE data increased this to 5818 classified SNVs (**Extended Data Table 6**). The successful classification of 5564 SNVs in *BRCA2* based on MAVE data, along with the new total of 5818 classified SNVs from MAVE, ClinVar and ENIGMA combined is expected to have important implications for the many carriers of these germline variants around the world. We note that the MAVE study accounted for 80.3% of all classifications, which represents a substantial impact on the VUS problem. Those with P or LP variants may now qualify for enhanced mammography and MRI screening and for surgical prevention to reduce the possibility of cancer through prophylactic mastectomy or oophorectomy. Furthermore, carriers may be eligible for treatment of breast, ovarian and potentially other cancers such as prostate and pancreatic with PARP inhibitors in the adjuvant and/or metastatic setting. In addition, family members of those with P or LP variants may benefit from testing and preventative measures and screening prior to onset of cancer. Moreover, those with B or LB variants can benefit from the knowledge that the variant that they carry is not a cancer predisposing allele.

Importantly, the MAVE thresholds were validated by three independent methods including ClinVar pathogenic and benign variants, orthogonal HDR assay functionally abnormal and normal variants, and nonsense and synonymous variants. Overall, the MAVE thresholds based on 95% prevalence of pathogenic and benign ClinVar standards yielded approximately 5% mis-categorization of the standards in each of the validation sets. While this raises the possibility of error in the ACMG/AMP/ClinGen clinical classification of BRCA2 SNVs, the need for multiple sources of evidence for a formal classification minimizes the likelihood of a misclassification error. However, as other MAVE, SGE or other functional assays are completed, consistency between the studies for each variant will be useful for overcoming any study specific errors. In a separate effort to further validate the MAVE findings, the MSK-IMPACT tumor sequencing dataset was used to assess whether functionally abnormal variants displayed loss of heterozygosity at the *BRCA2* locus, as should be observed for a driver mutation. Indeed, 86% of abnormal variants identified in MSK-IMPACT showed *BRCA2* LOH, whereas only 26% of functionally normal variants had corresponding LOH, suggesting a strong enrichment for loss of the wildtype second *BRCA2* allele in the tumors with functionally abnormal SNVs.

Case-control association analysis was also used to confirm that functionally abnormal SNVs were associated with an increased risk of breast and ovarian cancer. When combined together, the abnormal missense SNVs were associated with increased risk of breast cancer in both clinical high-risk and population-based studies and an increased risk of ovarian cancer in a clinical high-risk population. While public reference controls were used for the clinical high-risk analysis, the consistency of the findings confirmed the increased risk of cancer associated with these variants. Importantly, in a secondary analysis restricted to African American women, functionally abnormal variants were also associated increased risk of breast cancer. While studies of other races and ethnicities were not possible due to limiting numbers, it seems reasonable to suggest that the functionally abnormal variants will confer increased risk in all populations. Interestingly, while not significantly different, it was noted that the abnormal missense SNVs had lower risks than nonsense variants for both breast and ovarian cancer. This attenuation suggests that the threshold for functionally abnormal variants captured a number of hypomorphic missense variants with lesser effects on function and lower risks of cancer, although the attenuation could in part result from the intrinsic error in the MAVE data. Further studies of the BRCA2 SNVs to identify true hypomorphic variants with lower risks, that may require modified approaches to risk counseling and patient management, are needed.

The MAVE study had a number of limitations. In particular, the use of 95% thresholds resulted in intrinsic error in the categorization of variants. This threshold was chosen because only 11% of the SNVs were categorized as I/U variants that cannot be used for classification, whereas a 99% threshold classified many more variants as I/U. variants. The difference in loss of variants to the I/U category was in part a result of a limiting number of known pathogenic and benign standards for calibration. Further studies and comparisons with other functional assay datasets are expected to resolve some of the I/U variants and to confirm that the results from haploid HAP1 cells mirror results from diploid cells. In addition, because of relatively poor performance of the exon 23 and 26 MAVE experiments, the variants in these exons were categorized separately using a more conservative model and were subsequently integrated with results from the other DBD encoding exons. This resulted in loss of a larger proportion of SNVs in exons 23/26 to the I/U category. As before, other MAVE studies may resolve the categorization of many of these variants. While RNA studies were not conducted as part of this study, a number of SNVs in canonical splice sires, intronic regions or with high spliceAI scores were shown to be functionally abnormal, suggesting that the variants result in aberrant RNA splicing and protein truncation. Further studies of these variants, that are beyond the scope of the current study, are expected to establish whether the effects are directly on the protein or through aberrant splicing.

In summary, SNVs in the BRCA2 exons encoding the BD mutation hotspot were exhaustively characterized for effects on BRCA2 activity using a cell survival assay. The production of functional maps for 97% of all SNVs allowed for separation of nucleotide/protein-level functional aberrations and when coupled with genetic and clinical sources of evidence led to clinical classification of almost 4000 individual variants. These data will provide useful in the future, through integration with other datasets, for characterization and classification of all variants in this region in individuals from all racial and ethnic backgrounds and for all BRCA2 associated forms of cancer.

## Acknowledgements

This study was funded in part by NIH grants R35CA253187, R01CA225662, and a Specialized Program of Research Excellence (SPORE) in Breast Cancer to Mayo Clinic (P50CA116201), and the Breast Cancer Research Foundation (BCRF). Other sources of support were the Women’s Cancer Program Pilot Project Award at Mayo Clinic (SY), Halt Cancer at X National Research Grant (SY), Paul Calabresi Award in Clinical Translation Research at Mayo Clinic (K12CA090628) (SY), Career Development Award from the Conquer Cancer Foundation (SY), Sarasota Innovation Fund/Moffitt Foundation (ANAM), a Precision Prevention BCRF award (ANAM), and the Sigrid Jusélius Foundation (MM).

## Ethics Statement

All pedigrees and data shown in this paper are provided with the explicit written consent of the patients following IRB approval.

## Author contributions

**HH and CH** performed all SGE experiments and co-wrote the paper. **JN, SNH, TR, MA RDG, YAT, NB and WC** designed and performed data analysis and co-wrote the paper. **MR, MM, CB and DM** analyzed the MSK-IMPACT tumor data. **PCML and ANAM** generated figures and cowrote the paper. **MdlH** conducted Splice analysis. **SY and SMD** performed clinical interpretation of functional results.

## Methods

### Cell line and reagents

HAP1 cell (Horizon discovery) were maintained in IMDM with 10% FBS and 1% Penicillin/Streptomycin. For haploidy sorting, 1X10^-7^ HAP1 cells were resuspend in 5ug/ml Hoechst 34580 (BD, 565877) and sorted at 4°C. HAP1 cells were transfected using Turbofectin 8.0 (Origene). All oligos and primers were synthesized by Integrated DNA Technologies.

### Generation of site-saturation mutagenesis libraries and Cas9/sgRNA plasmids

BRCA2 DBD exons 15-26 and adjacent upstream and downstream 10 nucleotide introns flanking each exon were selected for saturation genome editing (SGE). Exons 18 and 25 were split into N-terminal and C-terminal targeted regions due to large exon size, resulting in 14 SGE target regions. Single guideRNAs (sgRNAs) were designed by Benchling design tool. sgRNA annealed oligos were ligated into pSpCas9(BB)-2A-Puro (PX459 v2.0) (Addgene; 62988) following BbsI (New England Biolabs, R0539L) digestion to create Cas9-sgRNA co-expression construct for each individual SGE. For each SGE, 600-1000bp homologous arms upstream and downstream of target region were amplified from WT HAP1 genomic DNA and cloned into BamHI-HF digested pUC19 vector using the NEBuilder HiFi DNA assembly Cloning Kit. Cloned plasmid backbones were subject to site-saturation mutagenesis by inverse PCR, ^30^ using mutagenized codon “NNN” primers for all possible nucleotide changes at each amino acid position. A Protospacer Protection Edit (PPE) encoding a synonymous mutation was introduced by site-directed mutagenesis into the protospacer adjacent motif (PAM) site or sgRNA recognition site of each target region to prevent re-cutting by the Cas9/sgRNA after successful editing. Furthermore, one 3-nucleotide mutation was introduced in the introns of each homologous arm to strengthen reamplification of the DNA in target regions.

### CRISPR/Cas9 saturation genome editing

Multiple sgRNAs with predicted high editing efficiencies on HAP1 cells were evaluated in SGE experiments of each target region and the optimal sgRNAs were selected (**Supplementary Table 1, 2**). In each SGE experiment, 2 million haploid-sorted HAP1 cells were seeded 24 hours prior to transfection, and co-transfected with 4μg target-specific variant library and 16μg Cas9/sgRNA targeting construct. Cells were selected in puromycin (1μg/ml) for 3 days. Cells were harvested at day 5 (24 hrs after puromycin selection) and day 14 post-transfection and genomic DNA (gDNA) was extracted using Monarch Genomic DNA Purification Kit (New England Biolabs, T3010L). Target regions were amplified by PCR to add barcodes for multiplexing. All PCR reactions were performed in 50 μL reactions using Q5 High-Fidelity 2X Master Mix (New England Biolabs, M0492L). Primers for genomic DNA amplification are included in Table S2. All reactions were cleaned and concentrated using Ampure XP beads prior to sequencing for 150 cycles on an Illumina MiSeq (approximately 5 million reads per run) or NextSeq (approximately 30 million reads per run). Base calls were performed by the instrument control software and further processed using customized algorithm.

### Sequencing data processing

FASTQ files of sequenced samples from Illumina MiSeq or NextSeq were trimmed for adapter sequences using cutadapt (v3.5). SeqPrep (v1.2) converted the paired-end reads into single reads. The single reads were aligned to the human reference genome (GRCh38) utilizing bwa-mem (v0.7.17). Following alignment, a custom-developed tool ‘CountReads’ was used for the analysis of DNA sequencing data, with a particular focus on mutation identification and characterization. ‘CountReads’ included the preparation of reference amino acid and DNA sequences, validation of sequencing data integrity, and precise trimming of reads to relevant regions. The method also differentiated between various variant types and confirmed the presence of specific variants and aggregated and reported variant data. ‘CountReads’ produced a VCF (Variant Call Format) file which was annotated with CAVA^31^. The SpliceAI tool (v1.3.1)^32^ was utilized to evaluate splicing effects associated for all observed single nucleotide variants (SNVs).

### Functional read count analysis

The log2 ratio between the frequency of day 14 and day 5 read counts was used to measure the depletion/enrichment effect for each variant. The comparison between experimental day 14 and day 5 avoided potential positional effect variation^15^. Variants with under-represented read counts (<10) in the library and day 5 were excluded from further analysis. Log2 ratios of variants were linearly scaled within each exon across replicate experiments relative to median synonymous and median nonsense SNV values. After within exon normalization, replicates of individual SNVs with Log2 ratios >1 relative to mean replicate values were excluded and Log2 ratios were recalculated. For each variant, the average score was calculated from all non-missing values among replicates. The linear scaling was used to normalize scores across exons using synonymous and nonsense SNVs, similarly as within exon normalization. Upper and lower bound cutoff scores for abnormal and normal variants were set by pathogenic and benign standards from ClinVar. Specifically, the thresholds of the function scores were set so that 95% precision for the abnormal group and the normal group, separated by an intermediate/uncertain (I/U) category was achieved. A total of 7013 DBD SNVs were evaluated after completion of all data cleaning and quality control (**Extended Data** Fig. 2).

### Three-dimensional structural modeling

BRCA2 functionally abnormal SNVs were mapped in the DBD using Pymol software. The PDB source file (PBD ID: 1MJE) was downloaded from the NCBI Molecular Modeling Database (MMDB). Three-dimensional structural modeling was based on the crystal structure of BRCA2-DSS1-SSDNA complex^33^.

### Multi-species amino acid sequence conservation and *In silico* pathogenicity prediction

BRCA2 amino acid sequences were obtained from Align-GVGD (http://agvgd.hci.utah.edu/).

Sequence alignments were performed using 10 species: *Homo sapiens, Pan troglodytes, Macaca mulatta, Rattus norvegicus, Canis familiaris, Bos taurus, Monodelphis domestica, Gallus gallus, Xenopus laevis,* and *Tetraodon nigroviridis*. Sequence conservation analyses were performed on amino acid residues containing BRCA2 DBD functionally abnormal variants. AlphaMissense^22^ and Bayes-Del^27^ were used for *in silico* pathogenicity prediction of BRCA2 DBD SNVs.

### Study populations

Breast and ovarian cancer cases and associated clinical phenotypes were collected from individuals receiving cancer genetic testing by Ambry Genetics. Public reference controls were non-cancer females from gnomAD (v4), excluding UK biobank. Breast cancer matching case-control data were also available from the CARRIERS and BRIDGES population based breast cancer studies^2,26^. Variants with allele frequency (AF)> 0.003 were excluded from the analyses.

### Comparison with other *BRCA2* functional assays

SGE functional results were compared with those from other studies, including a BRCA2-deficient cell-based homology-directed repair (HDR) assay^3,13^, a BRCA2-deficient cell line– based drug assay^23^, a prime editing–based SGE study^17^, and a mouse embryonic stem cell (m-ESC)–based functional analysis^24^.

### ACMG/AMP framework for classification of *BRCA2* DBD variants

The ACMG/AMP rule-based framework combines evidence from population, computational and predictive, segregation, functional, and other data, with each contributing source weighted as very strong (PVS1), strong (PS1, PS2, PS3, PS4), moderate (PM1, PM2, PM3, PM4, PM5, PM6), and supporting (PP1, PP2, PP3, PP4, PP5) evidence for pathogenic effects, or stand-alone (BA1), strong (BS1, BS2, BS3, BS4), and supporting (BP1, BP2, BP3, BP4, BP5, BP6, BP7) for benign effects. The combined data yield variant classifications of benign (B), likely benign (LB), pathogenic (P), likely pathogenic (LP), and variant of uncertain significance (VUS)^10^. In this study, ACMG/AMP scoring rules established by the ClinGen *BRCA1/2* Variant Curation Expert Panel (VCEP) were used for clinical classification of *BRCA2* DBD SNVs. The study was approved by the Western Institutional Review Board, which exempted review of the clinical testing cohort, and by the Mayo Clinic IRB. Detailed ACMG/AMP criteria used in this study can be found in supplementary materials.

### Tumor loss of heterozygosity (LoH) analysis

Loss of heterozygosity (LoH) status for breast, ovarian, pancreatic, and prostate cancer tumors carrying germine *BRCA2* DBD variants was acquired from tumor-normal paired sequencing using the Memorial Sloan Kettering – Integrated Mutation Profiling of Actionable Cancer Targets (MSK-IMPACT) panel^29^. The FACETS algorithm^34^ was used to determine LOH from matched tumor-normal pairs. Only tumor samples with >40% tumor content were included in the analysis.

## Statistical analysis

Associations between variant classification groups in *BRCA2* and the risk of breast cancer or ovarian cancer were performed using Fisher’s Exact tests using allele counts from Ambry Genetics female cases and gnomAD non-cancer females and for CARRIERS and BRIDGES matched breast cancer cases and unaffected female controls. Phenotypic comparisons between cases carrying functionally abnormal and normal variants were conducted using Student’s t-test for quantitative variables and Chi-Squared for qualitative variables. Lifetime absolute risks of breast cancer or ovarian cancer (malignant epithelial tumors of the ovary and fallopian tube) to age 80 years were estimated for different classification groups by incorporating odds ratio estimates with age-specific breast cancer/ovarian cancer incidence rates (restricted to non-Hispanic Whites) from the Surveillance, Epidemiology, and End Results (SEER) Program of the National Cancer Institute, accounting for all-cause mortality rates^2^. One-way ANOVA tests were conducted to compare the functional score differences of functional categories from other BRCA2 functional assays. Pairwise Wilcoxon rank-sum tests were performed to assess differences of *in silico* prediction scores in different MAVE functional categories. Fisher’s Exact tests were used in tumor LOH analysis. All analyses were performed with R software (v4.2.2) and all tests were two-sided. SGE data in bar graphs or scatter plots were presented as means from replicated experiments.

## Data availability

All data are presented in the article and/or in the supplementary materials and are available directly from the authors. All raw sequencing data were deposited into GEO, all related code was deposited in Github (<u>GitHub – najiemayo/Couch_SGE_BRCA2</u>).

