## Extended Figures and Tables for "Saturation genome editing-based functional evaluation and clinical classification of BRCA2 single nucleotide variants"

### Extended Data

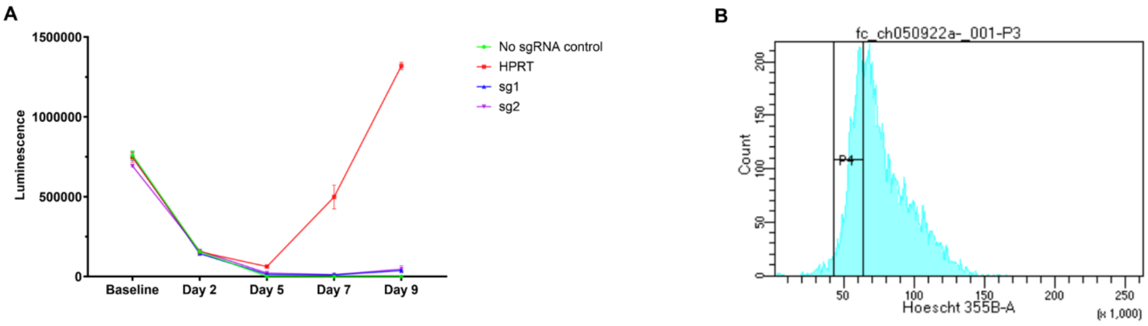


**Extended Data Fig. 1. HAP1 cell line essentiality confirmation and optimization for SGE.** **A**: Cell viability of HAP1 cells with no treatment control, Cas9/gRNA plasmid targeting non-essential gene HPRT, and two candidate Cas9/gRNA plasmids targeting exon 19 of *BRCA2*. Luminescence measured by the CellTiterGlow assay represented surviving cells upon targeting. Triplicate experiments were performed at each time point for each condition, and the luminescence counts were averaged. **B**: For maintaining haploidy of HAP1 cells in SGE experiments, cells were stained with Hoechst 34580 and subsequently sorted by a calibrated gate (P4) for selection of 1n HAP1 cells. sg: single guide RNA.

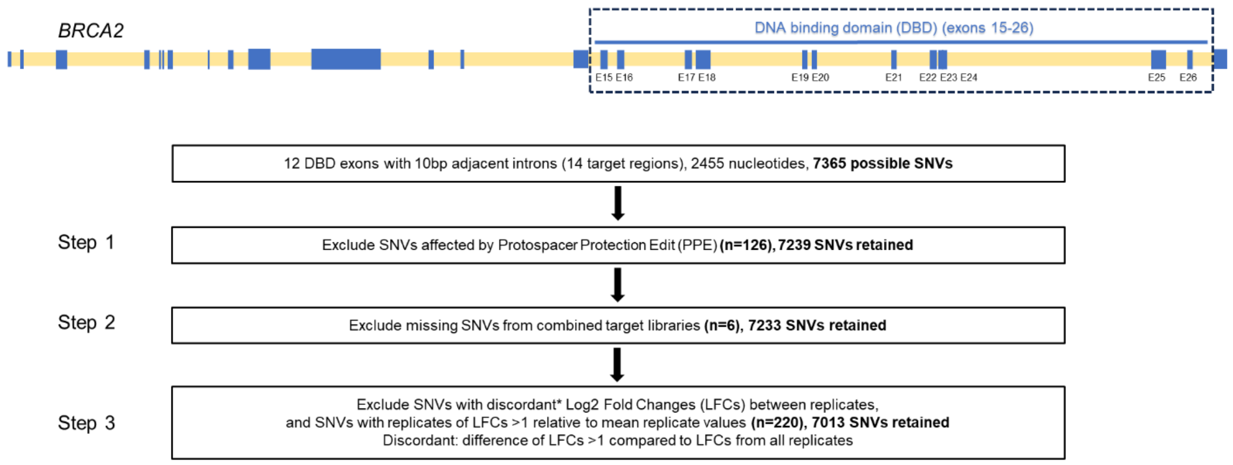


**Extended Data Fig. 2. SNV filtering strategies for accurate functional categorization.** The flow chart illustrates the filtering steps in the data processing and shows the number of SNVs excluded at each step.

**Extended Data Table 1. SNV recovery rate in the SGE experiments**.

| Target regions (include +/- 10 intron) | All possible SNVs | All possible missense SNVs | All possible nonsense SNVs | All possible synonymous SNVs | All possible canonical splice SNVs | All possible intronic SNVs | SNVs with functional score | Missense SNVs with functional score | Nonsense SNVs with functional score | Synonymous SNVs with functional score | Canonical splice SNVs with functional score | Intronic SNVs with functional score |
|---|---|---|---|---|---|---|---|---|---|---|---|---|
| E15 | 597 | 386 | 22 | 129 | 12 | 48 | 592 | 381 | 22 | 129 | 12 | 48 |
| E16 | 621 | 419 | 33 | 109 | 12 | 48 | 613 | 412 | 33 | 109 | 12 | 47 |
| E17 | 564 | 370 | 27 | 107 | 12 | 48 | 560 | 368 | 25 | 107 | 12 | 48 |
| E18N | 555 | 395 | 18 | 112 | 6 | 24 | 420 | 296 | 12 | 92 | 6 | 14 |
| E18C | 552 | 369 | 25 | 128 | 6 | 24 | 499 | 329 | 20 | 125 | 4 | 21 |
| E19 | 519 | 329 | 20 | 110 | 12 | 48 | 515 | 327 | 19 | 109 | 12 | 48 |
| E20 | 486 | 314 | 36 | 76 | 12 | 48 | 486 | 314 | 36 | 76 | 12 | 48 |
| E21 | 417 | 251 | 20 | 86 | 12 | 48 | 414 | 248 | 20 | 86 | 12 | 48 |
| E22 | 648 | 440 | 43 | 105 | 12 | 48 | 648 | 440 | 43 | 105 | 12 | 48 |
| E23 | 543 | 336 | 38 | 109 | 12 | 48 | 540 | 334 | 38 | 108 | 12 | 48 |
| E24 | 468 | 301 | 20 | 87 | 12 | 48 | 460 | 295 | 19 | 87 | 12 | 47 |
| E25N | 387 | 265 | 19 | 73 | 6 | 24 | 386 | 264 | 19 | 73 | 6 | 24 |
| E25C | 390 | 267 | 17 | 76 | 6 | 24 | 389 | 266 | 17 | 76 | 6 | 24 |
| E26 | 492 | 321 | 16 | 95 | 12 | 48 | 491 | 320 | 16 | 95 | 12 | 48 |
| Total | 7239 | 4763 | 354 | 1402 | 144 | 576 | 7013 | 4594 | 339 | 1377 | 142 | 561 |
| Percentages (%) | | | | | | | 96.88 | 96.45 | 95.76 | 98.22 | 98.61 | 97.40 |

Note: Variants in PAM/ /Protospacer Protection Edit (PPE) sites were excluded since they were not SNVs.
SNV: single nucleotide variant.

**Extended Data Table 2: MAVE functional prediction by variant type in each targeted region**.

| | variant type | Target regions (includes +/-10 intron) | | | | | | | | | | | | | | Total |
|---|---|---|---|---|---|---|---|---|---|---|---|---|---|---|---|---|
| | | E15 | E16 | E17 | E18N | E18C | E19 | E20 | E21 | E22 | E23 | E24 | E25N | E25C | E26 | |
| Functional abnormal | Canonical splice | 4 (33.3%) | 10 (83.3%) | 8 (66.7%) | 6 (100%) | 3 (75%) | 12 (100%) | 12 (100%) | 10 (83.3%) | 11 (91.7%) | 9 (75%) | 11 (91.7%) | 6 (100%) | 4 (66.7%) | 3 (25%) | 109 (76.8%) |
| | Intronic | 1 (2.1%) | 3 (6.4%) | 11 (22.9%) | 2 (14.3%) | 3 (14.3%) | 8 (16.7%) | 5 (10.4%) | 5 (10.4%) | 6 (12.5%) | 6 (12.5%) | 1 (2.1%) | 0 (0%) | 0 (0%) | 9 (18.8%) | 60 (10.7%) |
| | Missense | 20 (5.3%) | 71 (17.2%) | 81 (22%) | 43 (14.5%) | 73 (22.2%) | 71 (21.7%) | 7 (2.2%) | 5 (2%) | 26 (5.9%) | 25 (7.5%) | 26 (8.8%) | 31 (11.7%) | 20 (7.5%) | 23 (7.2%) | 522 (11.4%) |
| | Nonsense | 12 (54.6%) | 24 (72.7%) | 19 (76%) | 8 (66.7%) | 14 (70%) | 14 (73.7%) | 28 (77.8%) | 13 (65%) | 30 (69.8%) | 19 (50%) | 14 (73.7%) | 14 (73.7%) | 12 (70.6%) | 7 (43.8%) | 228 (67.3%) |
| | Synonymous | 3 (2.3%) | 5 (4.6%) | 1 (0.9%) | 3 (3.3%) | 2 (1.6%) | 6 (5.5%) | 1 (1.3%) | 3 (3.5%) | 2 (1.9%) | 4 (3.7%) | 2 (2.3%) | 0 (0%) | 2 (2.6%) | 2 (2.1%) | 36 (2.6%) |
| Functional I/U* | Canonical splice | 5 (41.7%) | 2 (16.7%) | 3 (25%) | 0 (0%) | 1 (25%) | 0 (0%) | 0 (0%) | 1 (8.3%) | 0 (0%) | 3 (25%) | 1 (8.3%) | 0 (0%) | 1 (16.7%) | 5 (41.7%) | 22 (15.5%) |
| | Intronic | 4 (8.3%) | 1 (2.1%) | 3 (6.3%) | 4 (28.6%) | 2 (9.5%) | 1 (2.1%) | 5 (10.4%) | 6 (12.5%) | 7 (14.6%) | 8 (16.7%) | 3 (6.4%) | 1 (4.2%) | 0 (0%) | 10 (20.8%) | 55 (9.8%) |
| | Missense | 52 (13.7%) | 60 (14.6%) | 44 (12%) | 43 (14.5%) | 44 (13.4%) | 38 (11.6%) | 16 (5.1%) | 19 (7.7%) | 57 (13%) | 43 (12.9%) | 16 (5.4%) | 28 (10.6%) | 20 (7.5%) | 68 (21.3%) | 548 (11.9%) |
| | Nonsense | 6 (27.3%) | 7 (21.2%) | 5 (20%) | 4 (33.3%) | 4 (20%) | 5 (26.3%) | 6 (16.7%) | 5 (25%) | 8 (18.6%) | 15 (39.5%) | 5 (26.3%) | 4 (21.1%) | 4 (23.5%) | 9 (56.3%) | 87 (25.7%) |
| | Synonymous | 10 (7.8%) | 4 (3.7%) | 2 (1.9%) | 12 (13%) | 11 (8.8%) | 11 (10.1%) | 1 (1.3%) | 7 (8.1%) | 10 (9.5%) | 12 (11.1%) | 2 (2.3%) | 12 (16.4%) | 6 (7.9) | 22 (23.2%) | 122 (8.9%) |
| Functional normal | Canonical splice | 3 (25%) | 0 (0%) | 1 (8.3%) | 0 (0%) | 0 (0%) | 0 (0%) | 0 (0%) | 1 (8.3%) | 1 (8.3%) | 0 (0%) | 0 (0%) | 0 (0%) | 1 (16.7%) | 4 (33.3%) | 11 (7.8%) |
| | Intronic | 43 (89.6%) | 43 (91.5%) | 34 (70.8%) | 8 (57.1%) | 16 (76.2%) | 39 (81.3%) | 38 (79.2%) | 37 (77.1%) | 35 (72.9%) | 34 (70.8%) | 43 (91.5%) | 23 (95.8%) | 24 (100%) | 29 (60.4%) | 446 (79.5%) |
| | Missense | 309 (81.1%) | 281 (68.2%) | 243 (66%) | 210 (71%) | 212 (64.4%) | 218 (66.7%) | 291 (92.7%) | 224 (90.3%) | 357 (81.1%) | 266 (79.6%) | 253 (85.8%) | 205 (77.7%) | 226 (85%) | 229 (71.6%) | 3524 (76.1%) |
| | Nonsense | 4 (18.2%) | 2 (6.1%) | 1 (4%) | 0 (0%) | 2 (10%) | 0 (0%) | 2 (5.6%) | 2 (10%) | 5 (11.6%) | 4 (10.5%) | 0 (0%) | 1 (5.3%) | 1 (5.9%) | 0 (0%) | 24 (7.1%) |
| | Synonymous | 116 (89.9%) | 100 (91.7%) | 104 (97.2%) | 77 (83.7%) | 112 (89.6%) | 92 (84.4%) | 74 (97.4%) | 76 (88.4%) | 93 (88.6%) | 92 (85.2%) | 83 (95.4%) | 61 (83.6%) | 68 (89.5%) | 71 (74.7%) | 1219 (88.5%) |
| Total variants | Canonical splice | 12 | 12 | 12 | 6 | 4 | 12 | 12 | 12 | 12 | 12 | 12 | 6 | 6 | 12 | 142 |
| | Intronic | 48 | 47 | 48 | 14 | 21 | 48 | 48 | 48 | 48 | 48 | 47 | 24 | 24 | 48 | 561 |
| | Missense | 381 | 412 | 368 | 296 | 329 | 327 | 314 | 248 | 440 | 334 | 295 | 264 | 266 | 320 | 4594 |
| | Nonsense | 22 | 33 | 25 | 12 | 20 | 19 | 36 | 20 | 43 | 38 | 19 | 19 | 17 | 16 | 339 |
| | Synonymous | 129 | 109 | 107 | 92 | 125 | 109 | 76 | 86 | 105 | 108 | 87 | 73 | 76 | 95 | 1377 |
| | All | 592 | 613 | 560 | 420 | 499 | 515 | 486 | 414 | 648 | 540 | 460 | 386 | 389 | 491 | 7013 |

I/U: intermediate/uncertain.

**Extended Data Table 3. Validation of the BRCA2 MAVE variant categorization thresholds by three independent datasets.**

| | | BRCA2 SGE functional categories | | | |
|---|---|---|---|---|---|
| | | Abnormal | Normal | I/U | Total |
| SGE internal controls | Nonsense | 182 | 19 | 59 | 260 |
| | Synonymous | 22 | 1030 | 82 | 1134 |
| Standardized HDR assay# | Abnormal | 87 | 6 | 13 | 106 |
| | Normal | 11 | 211 | 41 | 263 |
| | hypomorphic | 3 | 1 | 5 | 9 |
| ENIGMA/ClinGen VCEP standards | Pathogenic | 9 | 0 | 1 | 10 |
| | Benign | 2 | 43 | 5 | 50 |

#: a standardized V-C8 BRCA2 (-/-) HDR assay (Hu et al. AJHG 2023)

Note: all variants observed to influence splicing or predicted to influence splicing by SpliceAI were excluded in the validation datasets.

MAVE: multiplex assays of variant effect; SGE: saturation genome editing; I/U: intermediate/uncertain; HDR: homology-directed repair; ENIGMA: Evidence-based Network for the Interpretation of Germline Mutant Alleles; VCEP: variant curation expert panel.

**Extended Data Table 4: BRCA2 DBD subdomain specific multi-species amino acid residue conservation and comparison with MAVE functional results.**

| | | | | Perfectly conserved residues[$] | | | Highly conserved residues[!] | | | Poorly conserved residues[#] | | |
|---|---|---|---|---|---|---|---|---|---|---|---|---|
| BRCA2 subdomain | Wildtype residues | Evaluated residues[*] | Abnormal SNVs | Perfectly conserved residues[$] | Residues with abnormal SNVs (%) | Abnormal SNVs (%) | Highly conserved residues[!] | Residues with abnormal SNVs (%) | Abnormal SNVs (%) | Poorly conserved residues[#] | Residues with abnormal SNVs (%) | Abnormal SNVs (%) |
| Helical | 187 | 184 | 186 | 77 | 50 (65%) | 141 (76%) | 52 | 17 (33%) | 31 (17%) | 55 | 9 (16%) | 14 (7%) |
| OB1 | 134 | 131 | 136 | 47 | 32 (68%) | 78 (57%) | 47 | 23 (49%) | 43 (32%) | 37 | 12 (32%) | 15 (11%) |
| OB2 | 111 | 110 | 85 | 37 | 21 (57%) | 42 (49%) | 28 | 10 (36%) | 17 (20%) | 45 | 17 (38%) | 26 (31%) |
| Tower | 124 | 121 | 15 | 27 | 2 (7%) | 5 (33%) | 49 | 4 (8%) | 6 (40%) | 45 | 4 (9%) | 4 (27%) |
| OB3 | 130 | 128 | 93 | 19 | 13 (68%) | 31 (33%) | 48 | 19 (40%) | 30 (32%) | 61 | 19 (31%) | 32 (35%) |
| Subdomain linker region | 52 | 50 | 7 | 6 | 3 (50%) | 3 (43%) | 6 | 0 (0%) | 0 (0%) | 38 | 4 (11%) | 4 (57%) |
| Total | 738 | 724 | 522 | 213 | 121 (57%) | 300 (58%) | 230 | 73 (32%) | 127 (24%) | 281 | 65 (23%) | 95 (18%) |

*: residues containing the PAM/Protospacer Protection Edit (PPE) were excluded.
$: Perfectly conserved residues were defined as 100% residue sequence similarity across 10 species (pufferfish to homo sapiens).
!: Highly conserved residues were defined as 80-90% residue sequence similarity across 10 species (pufferfish to homo sapiens).
#: Poorly conserved residues were defined as ≤ 70% residue sequence similarity across 10 species (pufferfish to homo sapiens).
SNV: single nucleotide variant.
In this analysis, SNV is defined as changes result in missense alterations.

**Extended Data Table 5: Association analysis of MAVE functionally assessed variants with breast cancer in Non-Finnish European and African American populations.**

| | Case | | Control | | OR (95% CI) | P-value |
|---|---|---|---|---|---|---|
| | Number of variants | Number tested | Number of variants | Number tested | | |
| **NFE breast cancer - clinical testing cohort*** | | | | | | |
| MAVE missense abnormal$ | 137 | 123850 | 18 | 81097 | 4.99 (3.05-8.29) | $1.25X10^{-13}$ |
| MAVE missense normal | 1542 | 56358 | 2067 | 81095 | 1.07 (1.00-1.15) | $5.15X10^{-2}$ |
| MAVE nonsense abnormal | 82 | 123850 | 12 | 81085 | 4.47 (2.44-8.32) | $5.5X10^{-8}$ |
| DBD Pathogenic missense standard^ | 61 | 123850 | 7 | 81103 | 5.71 (2.67-12.63) | $3.96X10^{-7}$ |
| DBD Benign missense standard^ | 1965 | 56358 | 2038 | 81089 | 1.39 (1.31-1.48) | $5.74X10^{-24}$ |
| DBD nonsense | 102 | 123850 | 15 | 81085 | 4.45 (2.55-8.06) | $1.11X10^{-9}$ |
| **AA breast cancer - clinical testing cohort*** | | | | | | |
| MAVE missense abnormal$ | 19 | 16722 | 5 | 20537 | 4.67 (1.73-13.08) | $2.34X10^{-3}$ |
| MAVE missense normal | 295 | 6697 | 880 | 20537 | 1.03 (0.9-1.18) | $6.8X10^{-1}$ |
| MAVE nonsense abnormal | 50 | 16722 | 7 | 20537 | 8.78 (4-19.64) | $6.03X10^{-11}$ |
| DBD Benign missense standard^ | 716 | 6697 | 1813 | 20537 | 1.22 (1.12-1.34) | $1.8X10^{-5}$ |
| DBD nonsense | 51 | 16722 | 7 | 20537 | 8.96 (4.09-20) | $4.64X10^{-11}$ |

*: Clinical testing cohort collected cases from breast and ovarian cancer patients subjected to cancer genetic testing by Ambry Genetics, and controls from gnomAD v4 non-cancer females, excluding UK Biobank.

$: MAVE missense variants excluded potential splicing variant predicted by SpliceAI, variants with conflicting category with ClinVar interpretation, and AF > 0.003 in any population.

^: Missense standards are from ENIGMA reference panel.

NFE: non-Finnish European; AA: African American; OR: odds ratio; CI: confidence interval; MAVE: multiplexed assay of variant effect; DBD: DNA Binding Domain.

**Extended Data Table 6: Summary of MAVE ACMG/AMP classification.**

| **DBD SNVs** | ClinVar+ ENIGMA currently | MAVE ACMG/AMP classification | ClinVar+ ENIGMA variants after applying MAVE ACMG/AMP | MAVE ACMG/AMP classified not in ClinVar or ENIGMA | Classified in MAVE data from all sources | ClinVar+ ENIGMA not in MAVE | Total classified |
|---|---|---|---|---|---|---|---|
| B/LB | 993 | 4864 | 1887 | 3156 | 4995 | 48 | 5043 |
| P/LP | 417 | 700 | 497 | 278 | 748 | 27 | 775 |
| VUS | 1387 | 1449 | 413 | 915 | 1270 | 58 | 1328 |
| Total | 2797 | 7013 | 2797 | 4349 | 7013 | 133 | 7146 |

| **DBD Missense** | ClinVar+ ENIGMA currently | MAVE ACMG/AMP classification | ClinVar+ ENIGMA variants after applying MAVE ACMG/AMP | MAVE ACMG/AMP classified not in ClinVar or ENIGMA | Classified in MAVE data from all sources | ClinVar+ ENIGMA not in MAVE | Total classified |
|---|---|---|---|---|---|---|---|
| B/LB | 316 | 3084 | 1158 | 2053 | 3193 | 18 | 3211 |
| P/LP | 102 | 220 | 162 | 105 | 254 | 13 | 267 |
| VUS | 1293 | 1290 | 391 | 814 | 1147 | 58 | 1205 |
| Total | 1711 | 4594 | 1711 | 2973 | 4594 | 89 | 4683 |

MAVE: multiplex assays of variant effect; ACMG/AMP: American College of Medical Genetics/ Association for Molecular Pathology; DBD: DNA binding domain; SNVs: single nucleotide variant; ENIGMA: Evidence-based Network for the Interpretation of Germline Mutant Alleles; B: benign; LB: likely benign; P: pathogenic; LP: likely pathogenic; VUS: variant of uncertain significance.
Note: all variants evaluated are restricted to BRCA2 DBD SNVs including adjacent intronic +/-10nt region of each exon.
